# Single nucleotide variation catalogue from clinical isolates mapped on tertiary and quaternary structures of ESX-1 related proteins reveals critical regions as putative Mtb therapeutic targets

**DOI:** 10.1101/2023.06.22.507254

**Authors:** Oren Tzfadia, Axel Siroy, Alexandra Vujkovic, Abril Gijsbers, Jihad Snobre, Roger Vargas, Wim Mulders, Conor J. Meehan, Maha Farhat, Peter J. Peters, Bouke C. de Jong, Raimond B.G. Ravelli

## Abstract

Proteins encoded by the ESX-1 genes of interests are essential for full virulence in all *Mycobacterium tuberculosis* complex (MTBc) lineages, the pathogens with the highest mortality worldwide. Identifying critical regions in these ESX-1 related proteins could provide preventive or therapeutic targets for MTB infection, the game changer needed for tuberculosis control. We analysed a compendium of whole genome sequences of clinical MTB isolates from all lineages from >32,000 patients and identified single nucleotide variations (SNV). When mutations corresponding to all nonsynonymous SNPs were mapped on the surface of known and AlphaFold-predicted ternary protein structures, fully conserved regions emerged. Some could be assigned to known quaternary structures, whereas others could be predicted to be involved in yet-to-be-discovered interactions. Some mutants had clonally expanded (found in >1% of the isolates): these were mostly located at the surface of globular domains, remote from known intra- and inter-molecular protein–protein interactions. Fully conserved intrinsically disordered regions (IDRs) of proteins were found, suggesting that these are crucial for the pathogenicity of the MTBc. Altogether, our findings provide an evolutionary structural perspective on MTB virulence and highlight fully conserved regions of proteins as attractive vaccine antigens and drug targets. Extending this approach to other pathogens can provide a novel critical resource for the development of innovative tools for pathogen control.

## Introduction

*Mycobacterium tuberculosis* causes tuberculosis (TB) in humans and other mammals. This remarkably monomorphic pathogen shares 99.9% nucleotide similarity and identical 16S rRNA in its larger *Mycobacterium tuberculosis* complex (MTBc), unlike diversity seen in other bacteria (Böddinghaus et al. 1990; Sreevatsan et al. 1997; Achtman and Wagner 2008; Wiens et al. 2018). In the past decades, extensive research has been done to clarify the precise virulence mechanisms of the MTBc. ESAT-6 (6-kDa early secretory antigenic target, also known as EsxA) was identified as the main virulence-determining secreted protein (Andersen et al. 1995; Brodin et al. 2004). The avirulence of the century-old vaccine strain *M. bovis* Calmette-Guérin (BCG) is explained by deletion of the chromosomal region (RD1) containing *esxA* (Mahairas et al. 1996). Deletion of RD1 from the MTBc caused decreased virulence similar to that of BCG *in vitro* (Lewis et al. 2003). *M. microtii* spontaneously lost a similar RD1 region. It was shown that ESAT-6 interacts with CFP-10 and that this secreted heterodimer is critical for MTBc virulence through its cytolytic activity (Wel et al. 2007; Xu et al. 2007; Houben et al. 2012; Tiwari et al. 2019).

RD1 contains genes of the ESAT-6 secretion system 1, ESX-1 (Mahairas et al. 1996; Tekaia et al. 1999). ESX-1 is a member of the type VII secretion systems (T7SS), and is essential for full virulence in all MTBc lineages (L1–L8) as well as in the closely related pathogenic *M. marinum, M. kansasi* (Jagielski et al. 2020), and *M*. leprae (PMID: 11597336). Some genes necessary for ESX-1 transport and its regulation have been found outside the RD1 locus. The ESX-1 genes of interest (GOI) include multiple genes across different loci, required for the building and the functioning of ESX-1, the transport of virulence factors, and their membrane lysis activity. The ESX-1 GOI encodes 34 proteins (ESX-1 related proteins) that can be divided into four functional categories: 7 substrates (products that are secreted during virulence), 6 core complex (proteins part of the secretion machinery), 8 regulators (transcription factors), and 14 peripherals (exact function yet to be determined). The structures of two related T7SS inner membrane core complexes of MTB have been elucidated: ESX-3 (Famelis et al. 2019) and ESX-5 (Bunduc et al. 2021). The 3D structure of a few of these proteins (in isolation or as part of a complex) have been elucidated and validated, whereas predictions of all individual 3D protein structures have become available through artificial intelligence (AI) techniques (Baek et al. 2021; Jumper et al. 2021). Using the predicted 3D structures of individual proteins and knowledge of homologous protein complexes and interacting interfaces one could propose models for the quaternary structures of known interactions within the set of 34 ESX-1 related proteins.

Despite their high genomic similarity, MTBc lineages differ meaningfully in the host immune response, host tropism, phenotypes, drug resistance, and transmissibility (Brosch et al. 2002; Brosch et al. 2007; Wirth et al. 2008; Peters et al. 2020). So far, most research on the ESX-1 machinery has focused on MTBc L2 and L4 because of their widespread geographic range and availability of laboratory-adapted reference strains such as H37Rv (L4) and HN878 (L2). Thus, our knowledge on the genomic differences and convergence in the ESX-1 genes across the MTBc is limited.

To obtain a deeper understanding of virulence across MTBc, we generated a single nucleotide variation catalogue (synonymous and non-synonymous SNPs) in the ESX-1 GOIs, for all MTBc lineages (L1–L8) of human importance, using whole genome sequencing (WGS) data from >32,000 publicly available clinical isolates. First, we provide evidence illustrating the variation tolerance of the ESX-1 GOI, confirming that it is the most SNV-dense group of genes within the MTB genome. Next, we to identified several genomic regions including variants that arose independently under positive selection as done by Vargas et al (2022). We then examined the amino-acid locations that bear abundant SNV-counts. In 34 ESX-1 genes, only 21 SNPs were found in more than 1% of the isolates: these are considered successful fully functional transmission events. None of these resulted from convergence but either due to opportunistic sampling of the dataset or occurring in ancestral lineages. The data also reveal which parts of the 34 proteins are fully conserved. We mapped all SNVs onto each of the known or predicted 3D structures and inspected its surfaces. Varying sizes of conserved regions were found, some proteins showed clear polarity (of SNPs distribution). We then zoomed in on some essential motifs (such as Walker motifs, secretion signals, SS-bonds), and found <0.001% of isolates bearing SNVs in those locations. Next, we analyzed known and predicted quaternary structures, correlated interaction interface with SNV distribution maps, and experimentally validated that certain mutations on the interaction interface which still permit complex formation. We highlight a diverse set of conserved protein surface regions and hypothesize new interaction partners for these. Finally, we scrutinized the intrinsically disordered regions (IDRs) within our set of proteins, found two long stretches that are fully conserved, and discuss their potential essential role in immunological recognition. Combined, our findings highlight new targets for interfering with MTBc virulence.

## Results

### Consolidating SNPs for ESX-1 GOI

A collection of 32,399 unique MTBc isolates, including clinical MTBc isolates (Vargas et al. 2022) L1–L6 (NCBI), L7 (Chiner-Oms et al. 2019), as well as *M. bovis* (unpublished data), was collated (Figure 1, Supplementary Figure 1). The clinical isolates from human TB patients were considered a ‘filter’ for fully functional virulence. For the ESX-1 GOI, we examined 34 genes encoding 34 proteins with a total of 11,167 amino acids, for which a total of 8616 had a SNV, including 2742 synonymous mutations (sSNPs) and 5874 non-synonymous mutations (nSNPs). Almost 40% of the encoded amino acids had at least one nSNP. Figure 1 (and Supplementary Figures 3-6) visualizes the SNVs mapped on the predicted AlphaFold structures for each of the 34 ESX-1 GOIs.

**Figure 1.**
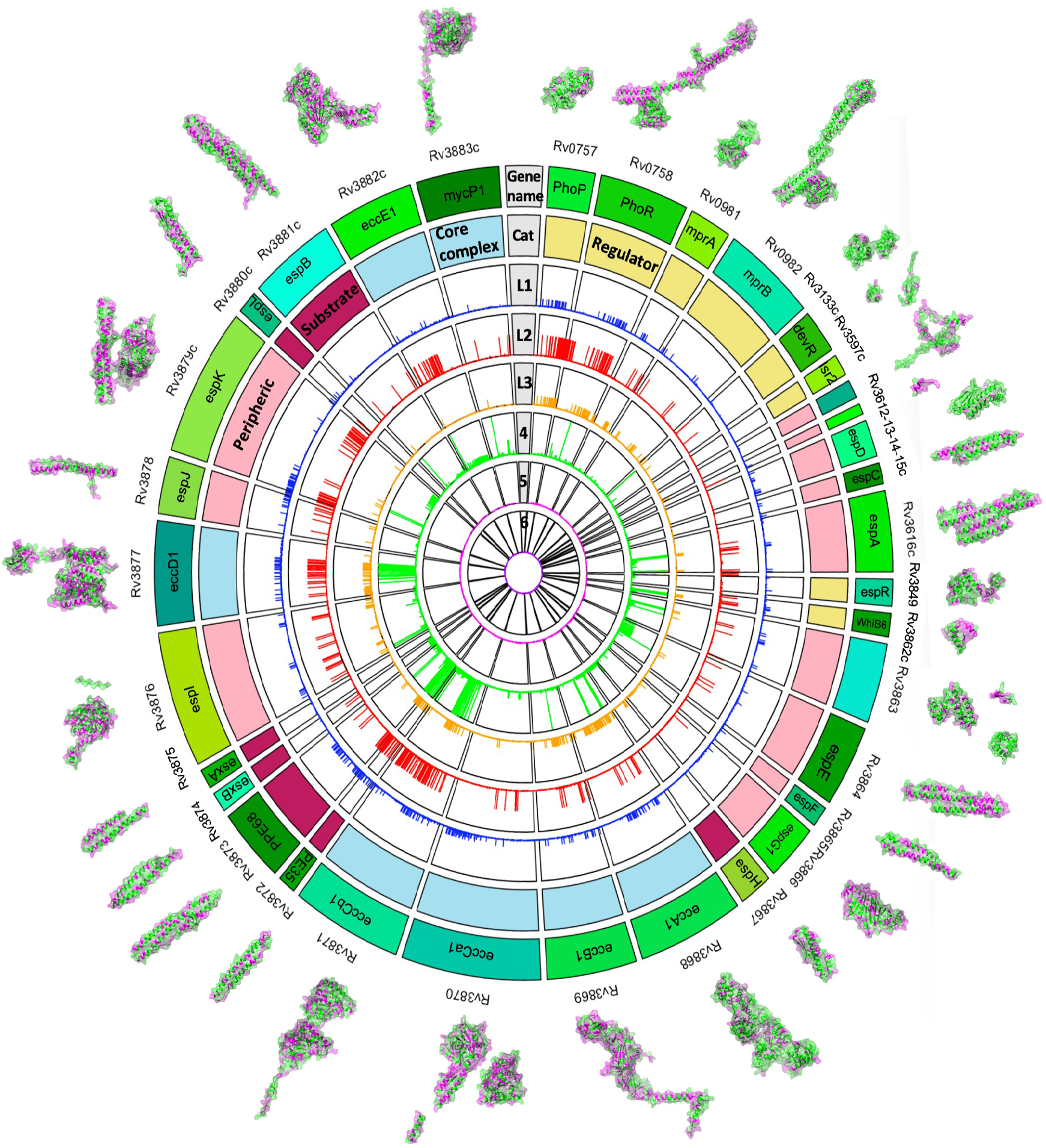
SNPs catalogue for 35 ESX-1 GOIs for each of the MTBc lineages. Circos plot visualization of distinctive nSNPs counted in the data was made from merged variant files (MTBseq pipeline) and then filtered by uniqueness per lineage (not shared with other lineages). The outer lane depicts gene names. Inner lane is color coded by functionality (purple – *substrates genes*, cream white for regulators, pale green for machinery coding genes, pale blue for peripheral coding genes. The next 7 inner lanes designated for lineage stratification (L1 – L6) (we exclude L9 due to small sample size (n=1)).

### Converging evolution

From convergence analysis, two silent SNPs in *espI* in four independent lineages (L1, L4, L3, and L8), in a total of ten unrelated clinical isolates were identified and confirmed by Sanger sequencing (Supplementary Table 2). EspI represses ESX-1 activity under conditions of lowered cellular ATP levels in the MTBc and may play a role during latent tuberculosis infection and reactivation (Zhang *et al*., 2015). In the isolates with two silent SNPs in espI, a repeat of 6 cytosines is formed. In cancer, this type of motif has been linked to changes in methylation (Dogan et al. 2017). Long PacBio reads could provide information about whether these mutations affect the methylation state of ESX-1. RNA-seq (gene expression profiling) on the mutant EspI strains followed by eQTL analysis will allow answering to what extent the SNPs in espI are biologically significant.

### Conservation on 3D level

We mapped all SNPs onto each of the known or predicted 3D structures and inspected the surfaces. Varying sizes of conserved regions were found, as shown by dominant green color of the predicted protein structures and the small number of red hotspots (Figure 2 and Supplementary Figures 3-6). Some proteins showed clear polarity, like WhiB6, PhoR, PhoP, DevR, MprA and MprB display SNPs on one side of their surface, while the opposite side showed no SNPs (Supplementary Figure 3).

**Figure 2.**
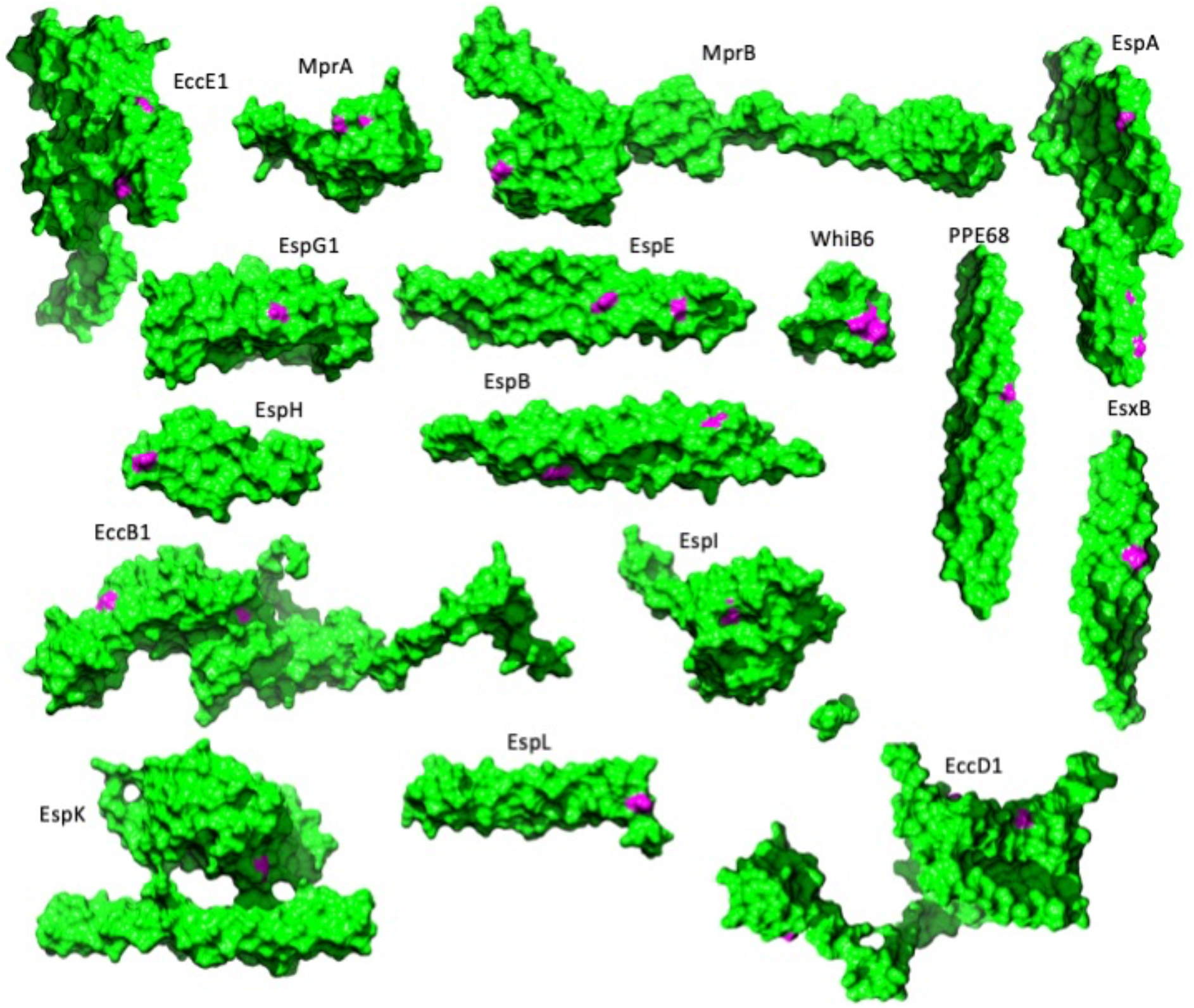
SNPs Hotspots. A hotspot is when the same mutation found in >300 clinical isolates (>1% of the isolates).

Comparing the prevalence of SNPs across the 34 ESX-1 GOI, we found that regulators WhiB6 (65.2%) and PhoR (54.2%) as well as two members of the ESX-1 inner membrane core complex EccB1 (54.4%) and EccE1 (53.8%) had the highest percentage of amino acid changes. In comparison, the regulator DevR was most conserved, with only 13.5% of its amino acids showing any mutation, and 98% of its SNPs were found at one position (see next section). This trend of clustered SNPs to one amino acid was also observed for *esxB* (96%), *PE34* (99%), *espA* (91%), *mprB* (94%), *mprA* (83%), *espH* (78%), *espE* (77%), and *espL* (69%) (Supplementary Figures 3–6).

### Hotspots / abundant SNPs in ESX-1 GOIs

After excluding polymorphisms due to convergent evolution, we identified 78% of the nSNPs in so-called hotspots (the same mutation found in >3000 clinical isolates, >1% of the isolates), reflecting a phylogenetically more ‘basal’ polymorphism with clonal expansion. We identified 21 hotspots (Figure 2), which involved 14 of the 34 ESX-1 GOI, involving all lineages (Supplementary Table 3). Only 4 proteins (PPE68, EccD1, EccE1, and EspK) showed more than 1 hotspot, while 9 (PhoR, MprA, MrpB, DevR, EspA, EccB1, EsxB, EspH, and EspB) each showed a single hotspot. Two other abundant SNPs were excluded as hotspots (T192I in EspA and E99* in PE34), since they included almost the entire dataset, suggesting that the H37Rv reference genome has a polymorphism, whereas one (L339H in MprB) included half the dataset. The hotspots occurred in all protein categories: we found them in 4 regulators, 3 peripheral, 4 substrates and 3 core components. Upon mapping the hotspots onto the protein structures, we found them predominantly on the surface of the protein, and to include both polar as well as apolar residues (Figure 2). Visual inspection of the experimental as well as predicted protein structures of these 14 proteins and 21 hotspots showed very few polar and H-bond interactions of the hotspot side chains within the protein itself (data not shown). Future identification of potential compensatory mutations may explain how these clonally expanded mutations are non-detrimental nor with other subunits for those proteins where experimental complex structures were available (more below). For example, in EsxB the most prevalent mutation was identified as E68K. Chimera predicted it to have minimal impact on the binding site between EsxB and EsxA. To further investigate this finding, we are currently cloning a mutant strain and assessing its protein interaction with EsxA.

### Motifs and binding regions

Next, we analysed the SNPs found for the two motifs known to be essential for secretion in the 7 substrates: WxG and YxxxD/E. Overall, these secretion motifs were found to be highly conserved. Within EsxB, only one SNP was found in WxG and two in YxxxD/E. Within EpsJ, the YxxxD/E motif had 11 total SNPs. For YxxxD/E, and EspH had 6 SNPs EspF had 1. The YxxxD motif is fully conserved in EspA and EspC.

Some of the ESX-1 core components contain ATPase domains. We analysed the Walker A motifs in EccA1, EccCa1 and EccCb1. The Walker A motif, also called the “P-loop” for its phosphate binding, has the classical pattern (G/A)xxxxGK(T/S) (Allemand, Maier, and Smith 2012). Our data show a total of 4 SNPs within this motif for the total of five ATPase domains found in EccCa1 and EccCb1, no SNP found in EccA1 motif (Supplementary Table 4). The Walker B motif (hhhhD, where h is any hydrophobic residue) is fully conserved or has allowed changes in all three proteins. We next analysed the conservation of disulfide bonds. EccB1 has a disulfide bond within its periplasmic domain (CysXX-CysYY): it is fully conserved. Cys48 in substrate EspC has been described to play a role in EspC polymerisation and subsequent dimer and disulfide bond formation (Lou et al. 2017). No mutations were found for this residue. Finally, we evaluated the Fe-S cluster in regulator whiB6, to which NO can bind as part of the innate immune response to mycobacterial infection. This binding inhibits the secretion of ESAT-6 and CFP10. We found no SNPs in the binding pocket of this Fe-S cluster. As for the inner membrane core complex ESX-1 genes, it appears that the protein tails of EccCb1, EccD1, MycP1, are all SNP free. We modelled the ESX-1 core complex as a trimer of dimers and colored all its components according to the number of SNPs. Zooming out on the complex, it seems to contain mutation-free pockets. As of today, there is no experimental structure known of this complex. Our dataset reveals a large overall number of mutations within each of its components. These might include mutation-pairs, in which interactions are maintained using different pairs of amino-acids. Detailed analysis of those awaits better experimental insight in its structure. Together, these data demonstrate that functional regions within the set of 34 ESX-1 GOI are largely conserved.

### Intrinsically disordered regions (IDRs)

The predicted AlphaFold structures contain regions of low and very low model confidence scores (respectively, <70 and <50). The percentage of low-confidence regions varies for different species, and is relatively small for bacterial proteomes such as *M. tuberculosis* (13.29%) (Aderinwale et al. 2022). We analysed whether these regions of low confidence in the proteins encoded by the ESX-1 GOIs reflect intrinsically disordered regions (IDRs) (Piovesan, Monzon, and Tosatto 2022), which tend to be hydrophilic unstructured protein structures thought to be disproportionally involved in interactions. Thirteen of the 34 ESX-1 GOI proteins contained at least one IDR, as defined both by low pLDDT confidence score and IDR-analysis schemes; Supplementary Figure 3). In particular, all seven substrates have large stretches of IDRs: EspA, EspE, PPE68, EspI, EspJ, EspB and EspK. Few nSNPs and no hotspots were found within these regions. We analysed the percentage of mutated amino acids within the IDRs of each of these proteins and found a strikingly low percentage within IDRs of EspE, EspI, EspK and EspB (Figure 3). EspB even displayed a long, entirely nSNP-free IDRbetween the structured N-terminal domain (residues 1–262) and the MycP cleavage site (residue 345); a second IDR follows that maturation site and contains a few nSNPs. MycP1 cleaves off this second IDR within the periplasm, after the translocation of EspB through the ESX-1 inner membrane core complex. Structural studies, including macromolecular crystallography and cryo-electron microscopy, could not reveal a structural insight about the first IDR (residues 262–375).

**Figure 3.**
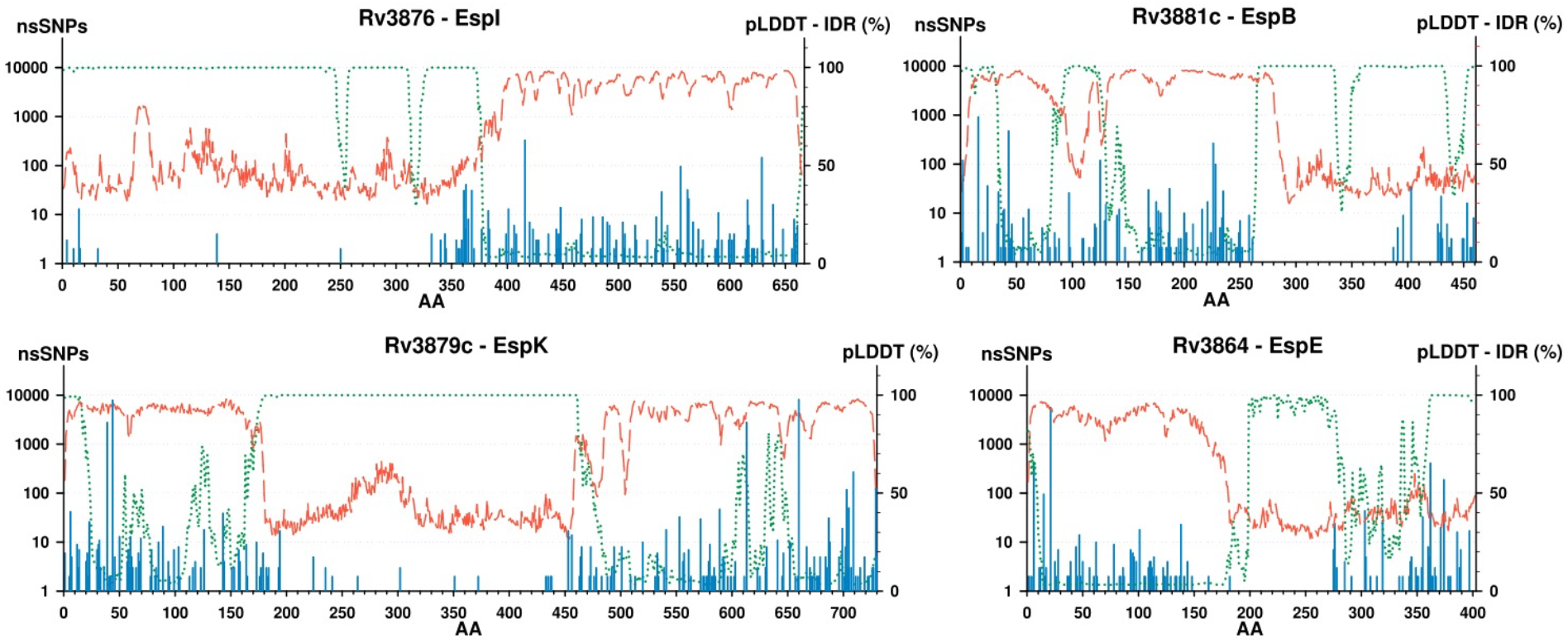
ESX-1 components with the most conserved predicted IDRs. The amino-acid sequences of the ESX-1 proteins of interest were analyzed with the AlphaFold2 and ODiNPred algorithms. The proteins with the longest low-pLDDT, high-disorder scores (IDR), low nSNP frequency stretches were identified as EspE (>100 AA), EspB (<150 AA), EspK (∼250 AA) and EspI (<350AA). Per amino-acid position: nSNPs counts, AlphaFold2 confidence and ODiNPred disorder scores were plotted with -respectively -vertical bars (blue, left axis, logarithmic scale), dashed red (pLDDT score) and green dots (IDR score) lines (right axis, 0-to-100% scale).

### Quaternary structures

Because the T7SS uses several protein complexes for the transport of virulence factors in pathogenic mycobacteria, we also inspected the nSNPs within the context of quaternary structures to obtain insight into the fidelity of these interfaces for interaction. Both known and putative complex interfaces were examined using the PISA interface tool (Krissinel and Henrick 2007). For each MTB complex interface, we determined the number of SNPs for each interacting residue. Our analysis revealed interesting variations in SNP counts among different protein-protein and protein-DNA interactions (Supplementary Tables 7 -12) (Figure 4). In addition to quaternary structure interactions, we determined the presence of SNPs in experimentally verified regions where several ESX-1 substrates interact.

**Figure 4.**
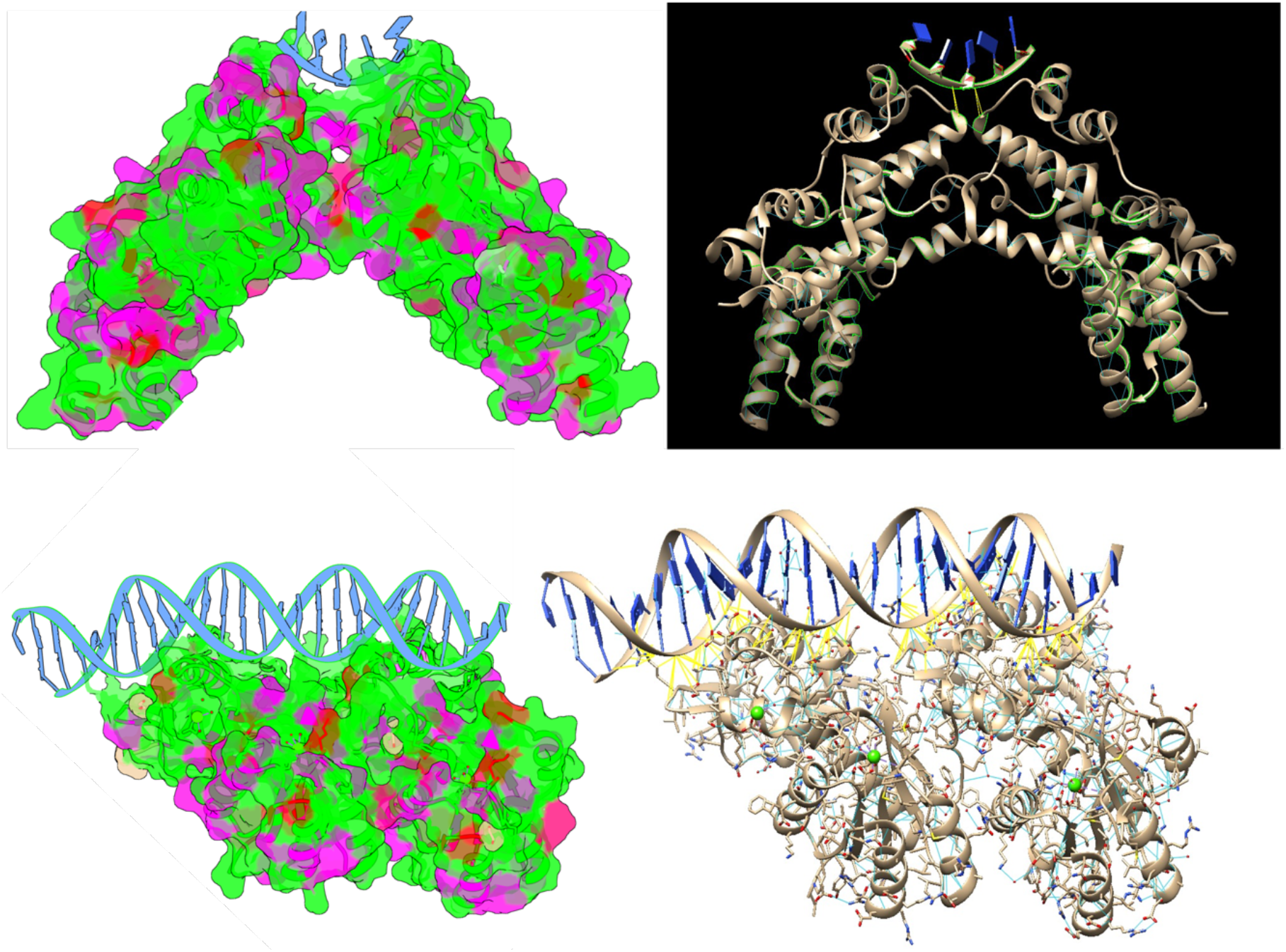
EspR (top) and PhoR (bottom) display interfaces with DNA clean of SNPs

The main substrates ejected from the phagosome by ESX-1 are EsxA and EsxB, known to form a dimer before being secreted. Both proteins have only 2 residues with 2 SNPs detected (Supplementary Table 7). Surprisingly, when analyzing the self-interaction of EspB and PhoR, where we count total of <50 SNPs (in >30k isolates), in <25% of interacting residues (Supplementary Tables 8-9). This observation suggests the presence of natural variations within the EspB and PhoR proteins, potentially indicating functional diversity among MTB strains. Further studies are needed to elucidate the impact of these variations on the overall virulence and pathogenicity of MTB.

Furthermore, in the interaction between EspK and EspB, we observed only 5 residues with SNPs count 1-15 SNPs. The other 10 residues known to interact had no SNPs detected in 32k isolates, suggesting a high degree of conservation in this interaction rendering stability and fidelity of the EspK-EspB complex (Supplementary Table 10).

Regarding the interaction between regulators such and EspR and PhoR with DNA, we see complete conservation in all 11 interaction sites (7 residues in PhoR – DNA and 4 residues in EspR and DNA; Supplementary Tables 11-12). The conservation of this interaction site indicates its functional importance for proper gene regulation and underscores its significance in the context of MTB’s virulence mechanisms.

## Discussion

These studies demonstrate that 34 genes encoding ESX-1 proteins were somewhat conserved. For example, *espF* had 44, *esxA* had 39, and *espC* had only 37 SNVs across their respective sequences. In cases where >50% of the amino acid positions had ≥1 SNP, we define it as *mutated protein*. Of the 34 ESX-1 GOIs analysed, only 6 mutated proteins were identified: EccB1, EccD1, EspJ, PhoR, Rv3613c and WhiB6.

When examining the distribution of SNPs across the ESX-1 GOI proteins, captivating patterns emerge, which can shed light on both known and unknown proteins’ functions and interactions. Mapping these onto experimental and predicted 3D structures revealed biased SNV distribution patterns. Mutation hotspots seem to occur mostly at places on the protein surface for which no interactions have been reported so far. The observed clustering of SNPs to a single amino acid was not surprising (Supplementary Figures 2–3), as these are known to be prone to polymorphisms. The hotspots suggest that these regions can vary without functional consequence, and thus have no evolutionary pressure to remain stable.

Interestingly, IDRs also harbored relatively few SNPs (Figure 3). This sequence conservation across 34 genes from >32,000 isolates indicates that these protein regions play crucial roles in the virulence of *M. tuberculosis*.

Conservation plays a crucial role in maintaining the integrity and functionality of proteins involved in the ESX-1 virulence mechanism of MTB. One important aspect of conservation is the absence of SNVs in the interaction sites of these proteins.

Most proteins have signature sequences or motifs that are characteristic of protein families, these motifs represent an important feature in the protein structure or function. ESX-1 is a molecular motor that pumps proteins through mycobacterial membranes using the chemical energy of ATP hydrolysis. The Walker A and B motifs are motifs founds in ATPases and GTPases involved in the nucleotide binding and hydrolysis. Any variation in the sequence could potentially affect any of these two capacities, for that reason, it is not surprising the conservation level observed in ESX-1 ATPases. It is noteworthy that even ATPase 2 and 3 domains, which correspond to EccCb1, are conserved even though it has been suggested that these domains are not catalytically active (ref). This idea is based on residues orientation in EccC structure from Thermomonospora curvata and the fact that the crystal structure included ATP molecules in these domains. However, it is possible that those observations do not represent what happens in all EccCb’s as we have evidence of ATPase activity in EccCb1 from *M. tuberculosis*). In all >32k samples, four variations in the Walker A glycines were found, either for an alanine or arginine. These residues are involved in the structural integrity of the motif or the coordination to the ATP β-phosphate, which could be affected with those changes; however, it is important to note that the actual impact of this mutation would depend on the specific protein, its three-dimensional structure, and its functional context. Experimental studies and structural analyses would be necessary to determine the precise consequences of such a mutation on the protein’s activity.

Further investigation is necessary to determine the functional consequences of these variations and their potential implications for MTB’s ability to manipulate host responses.

Overall, our analysis of SNP counts in specific interactions within the ESX-1 virulence mechanism of MTB provides valuable insights into the conservation and genetic diversity within these critical protein-protein and protein-DNA interfaces. These findings contribute to our understanding of the functional significance of these interactions and their potential implications for MTB’s pathogenicity. Many proteins contain signature sequences (motifs) that are characteristics of a protein family. These signature sequences are part of important structural or functional domains.

## Methods

### Whole genome sequencing (WSG) and phylogenetic analysis of 971 clinical isolates

A total of 971 genomes of clinical isolates from the MTBc (*n*=965; L1–L8) and from *M. bovis* (*n*=6) were included, from Bangladesh and Gambia (Comas *et al*., 2013; Lempens *et al*., 2020; Ngabonziza *et al*., 2020). We used this data set as the initial backbone phylogeny for the rest of the analysis. The semi-automated MTBseq pipeline was used for reads mapping and variant detection (Kohl *et al*., 2018). The output of MTBseq was used to generate an SNP alignment using an in-house Python script (https://github.com/alxndravc/ESX-1-MS). Based on this SNP alignment, a maximum likelihood tree was built using RaxML-NG (Stamatakis, 2014) with a GTR + CAT model of evolution and 100 bootstraps; *M. canetti* was added as an outgroup. The different phylogenetic lineages were visualised using the online interactive Tree Of Life (iTOL) tool (Letunic and Bork, 2019).

### Phylogenetic analysis of 31,428 publicly available MTBc isolates

We used a SNP barcode (Freschi *et al*., 2020) to type a collection of 31,428 WGS MTBc isolates downloaded from NCBI into MTBc lineage and sub-lineage. We excluded isolates that were missing 10% of SNP sites, were not typed as belonging to MTBc L1–8 or were typed as L4 but were not typed further with an L4 sub-lineage. We split the 31,428 isolates into eight groups based on genetic similarity, five groups corresponding to global L1, L2, L3, L5, L6 and three groups for lineage 4 (i.e., L4.1.12). To generate phylogenies for each of these groups, we first merged VCF files of the isolates in each group with bcftools (Li *et al*., 2009). We then removed repetitive, antibiotic resistance and low-coverage regions (Freschi *et al*., 2020). We generated a multi-sequence FASTA alignment from the merged VCF file with vcf2phylip (version 1.5). Finally, we constructed the phylogenetic trees for each group with IQ-TREE 1.6.12 (Nguyen *et al*., 2015). We used the *mset* option to restrict model selection to GTR + CAT models and selected the GTR+F+I+R model for the six isolate groups corresponding to L1–L4 and implemented the automatic model selection with ModelFinder Plus (Kalyaanamoorthy *et al*., 2017) for the isolate groups corresponding to L5 and L6.

### SNP catalog

An in-house Python script was used to count the unique SNPs present within each isolate group (stratified by global lineage) within the ESX-1 GOIs (https://github.com/alxndravc/ESX-1-MS). To calculate the SNP frequency per 1 kb, the number of unique SNP locations per gene were multiplied by 1000 bp and divided by their respective gene length. The same methodology was used on the 31,000 WGS isolates from NCBI. The genes were divided into 4 groups: machinery, substrates, regulatory and peripheral (i.e. genes not belonging to any of the other three categories) (Supplementary Table 1). One-way ANOVA with posthoc Tukey was used to determine between which pairs of means there is a significant difference (P < 0.05), excluding the genes in the unknown group. Z-scores were determined as standard deviations from the mean for each of the genes, with a cut-off set at 1.5 to indicate genes with significant SNPs. To count the SNPs that represented at least 20% of the total isolates per lineage, SAMtools and IGV were used. The IGV coverage allele-fraction threshold was set at 0.2, i.e., if a nucleotide differs from the reference sequence in greater than 20% of reads, IGV colors the bar in the coverage bar chart in proportion to the read count of each base (A, C, G, T).

AlphaFold2-generated protein 3D models were collected from the EBI/AlphaFold collection of models built upon the UniProt database (version 4, last accessed in 20/7/2022, available at https://alphafold.ebi.ac.uk), using their corresponding UniProt reference. For each model, pLDDT scores per amino-acid position were extracted from the PDB files (b-factor column), and nSNP counts per amino acid were assigned into corresponding attribute files prior to their rendering with UCSF Chimera 1.15 and/or ChimeraX 1.4 (ref, websites). Singletons were filtered out by setting up the following color oding scheme: green for 0 or 1 nSNP, purple for a minimum of 2 nSNPs, red for the 1% outliers (>300 nSNPs).

### nSNPs vs pLDDT vs IDR plots

Putative Intrinsic Disordered Regions (IDRs) of the 34 ESX-1 Proteins of Interest (POIs) were predicted in batch using the OdiNPred server (Prediction of Order and Disorder by evaluation of NMR data, https://st-protein.chem.au.dk/odinpred) without evolution and the Disorder Probability (DP) scores were retained for further processing. nSNP counts, pLDDT scores and DP scores were assigned to each amino acid position, per POI, and DP scores were normalized to % for plotting purpose. Figures were generated with SigmaPlot 12.5 (Systat Software): nSNPs counts per AA were displayed on a logarithmic scale (1 to 40000) to visually filter singletons out (left axis) while pLDDT and DP scores per AA were displayed on a 0-to-100% scale, (right axis).

### SNPPar analysis homoplasy

The resulting phylogenetic tree from the 971 clinical isolates was used in SNPPar along with the SNP dataset to obtain the mutation events across all MTBc lineages. To screen for convergent SNP sites in the alignment, SNPPar was used. Based on the provided phylogenetic tree, SNPPar searches for SNPs that are the same mutation (e.g., C G) at the same position in two or more unrelated isolates or different mutations that result in the same base (e.g., C G, A G) on the same position. It also detects revertant mutation back to the ancestral state (e.g., C G C) (Edwards *et al*., 2020). We used the default settings of SNPPar, which is a TreeTime for ancestral state reconstruction (ASR). As input, the phylogenies mentioned above were used together with the H37Rv reference genome in Genbank format (NC_000962.3) and an SNP position file. On the large dataset of 31,428 publicly available samples, SNPPar was run 8 times on 8 independent sets of isolates corresponding to 8 genetic backgrounds (L1–L6). Lineage sample counts are as follows: L1: 2,815; L2: 8,090; L3: 3,398; L4A: 5,839; L4B: 6,958; L4C: 4,134; L5: 98; and L6: 96.

### Quarterly structure interfaces analysis

Understanding the molecular interactions and stability of complexes in Mycobacterium tuberculosis (MTB) is crucial for deciphering its pathogenesis and identifying potential therapeutic targets. In this study, we provide a detailed analysis of experimentally verified MTB complexes, focusing on their interface characteristics, energy stability, and the presence of single nucleotide polymorphisms (SNPs) (Supplementary Tables 7-12).

### Sanger sequencing of espI

To validate the two *espI* silent SNPs (Pro134/135Pro), primers were designed for Sanger sequencing on the DNA extracts of available mutant isolates (i.e., containing the SNPs) and phylogenetically closely related wild-type (WT) isolates. A list of used genetic isolates and primer sequences can be found in Supplementary Table 2.

